# A minority-ruled population coding of kinematics in the striatum

**DOI:** 10.1101/130237

**Authors:** Wahiba Taouali, Pavel E. Rueda-Orozco, David Robbe

## Abstract

The structure of population activity in the dorsolateral striatum during the performance of motor sequences has not been characterized and it is unclear if striatal ensembles encode (predict) kinematic parameters defining how sequences are executed. Here we analyzed hundreds of striatal spike trains from naive and trained rats performing a running sequence. We found that the population response was composed of a diversity of phasic modulations covering the entire sequence. The accuracy of kinematics encoding by single neurons was around chance level but improved when neuronal ensembles were considered. The distribution of single-neuron contributions to ensemble encoding was highly skewed with a minority of neurons responsible for most of the encoding accuracy. Importantly, running speed ensemble encoding improved after learning. We propose that during motor learning, striatal ensembles adjust their task representation by tuning the activity of a minority of neurons to the kinematic parameters most relevant to motor performance.

## Introduction

There are converging evidences from a large body of experimental and theoretical work that the basal ganglia control how actions are performed (Shadmehr and Krakauer, 2008; Turner and Desmurget, 2010; Dudman and Krakauer, 2016). Several studies have pointed to the dorsal region of the striatum, the main input nucleus of the basal ganglia, as a key determinant of movements vigor that is their speed, amplitude and frequency. Parkinsonian patients, whose dorsal striatum is severely depleted in dopamine, are still capable of executing accurate reach movements but systematically use lower speeds than healthy control subjects, suggesting a specific contribution of this brain region to the selection of movement speed (Mazzoni et al., 2007). In mice, perturbing activity of dorsal striatal neurons reduced the speed of forelimb reaching movements to displace a joystick (Panigrahi et al., 2015) and selectively impair state-modulation of response vigor in a nose-poking decision task (Wang et al., 2013). Putative neuronal correlates of movement speed control have been reported in the dorsal striatum. First, electrophysiological recordings performed in different subregions of the dorsolateral striatum (DLS) revealed perfect linear correlations between the firing rate of neurons and the movement speed of different body parts such as the forelimbs, neck, whiskers and tong (West et al., 1990; Carelli et al., 1997; Pederson et al., 1997; Tang et al., 2007; Kim et al., 2014). While these activities might contribute to movement speed control, it can not be ruled out that they reflect passive somatosensory responses to joint rotation (Mink, 1996). Second, moderate correlations between striatal neurons firing rate and task-related kinematic parameters (including speed) have been observed in mice performing reaching movements (Panigrahi et al., 2015) and rats performing a fine-tuned running sequence (Rueda-Orozco and Robbe, 2015). The correlation coefficients rarely exceeded 0.5, ruling out passive sensorimotor responses as their main explanation. However, it is not clear if these relatively noisy correlations can support a representation of speed (or of other kinematic parameters defining how the sequence is performed), and by representation we mean the capacity to predict, at different phases of the sequence performance, the value of kinematic parameters from only the spiking activity of single neurons or neuronal ensembles.

To address this question, we took advantage of recordings of spiking activity of dorsolateral striatal neurons from naive and trained rats performing a well-defined running sequence on a motorized treadmill (Rueda-Orozco and Robbe, 2015) and 1) characterized at the population level the variability of striatal spiking responses during sequence performance, 2) applied a non-parametric method to quantify, beyond linear correlations, the dependence between task-relevant kinematics parameters (running speed, acceleration and position of the animals on the treadmill) and the response of striatal neurons and 3) evaluated the decoding of kinematics parameters by individual neurons and neuronal ensembles using the Bayesian inference framework. Our analyses revealed that the striatal population activity is 1) composed of a diversity of heterogeneous phasic modulations covering the entire task performance, 2) predictive of the animals running speed and position on the treadmill, with a minority of strong contributors responsible for most of the prediction accuracy and 3) selectively tuned by motor learning, as running speed population decoding was more accurate in trained versus naive animals.

## Results

The structure of population activity in the dorsolateral striatum during motor sequences has not been determined and it is unclear if striatal ensemble can encode parameters defining how sequences are executed. To address these issues we analyzed the pattern of activity of hundreds of striatal neurons recorded while rats performed a stereotyped running sequence on a motorized treadmill. The sequence was composed of three elements: passive displacement from the front to the rear portion of the treadmill, stable running around the rear portion of the treadmill and acceleration across the treadmill to enter the reward area. The animals were either self-trained (ST, i.e., they had learned this sequence through a long trial-and-error process) or were naive to the task and hand-guided (HG) to perform the running sequence. The recording sessions were composed of trials, during which rats perform the motor sequence, interleaved with intertrial resting periods (**Figure 1A**, top panel, (Rueda-Orozco and Robbe, 2015)).

**Figure 1:**
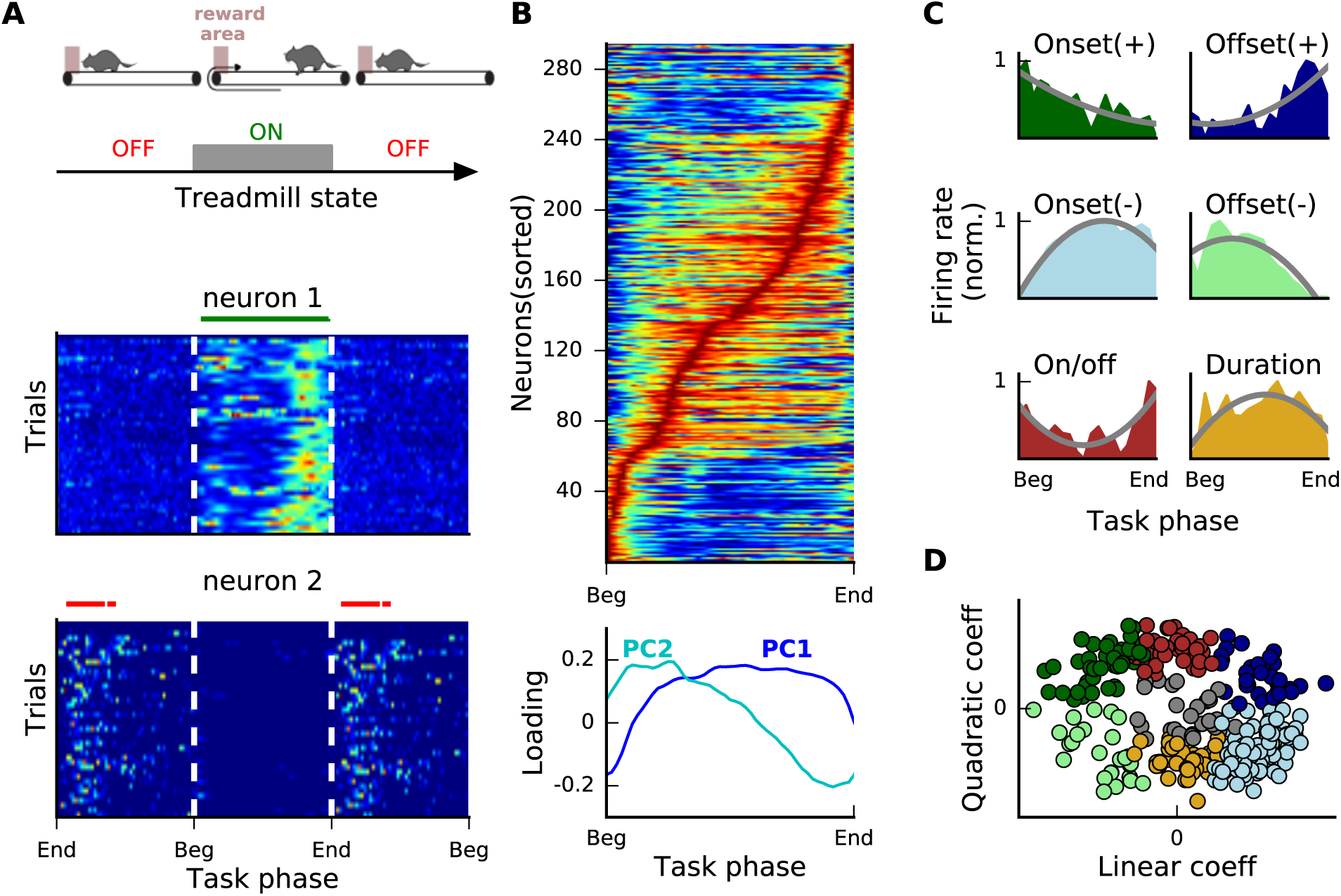
A continuous and heterogeneous representation of the running sequence in the striatum. (A) Top: Schematic representation of the task. Middle and Bottom: Trial-by-trial color coded firing rate (blue for min and red for max) of two example neurons showing an increase of their activation during the trial (neuron 1) or the intertrial (neuron 2) period. Horizontal bars show the phases with significant modulations (Methods). (B) Top: Normalized average firing rates (sorted according to the task phase of the maximum firing rate) of positively modulated neurons recorded in three ST animals. Same color code as in (A). Bottom: First two principal components (PC1: dark blue, PC2: light blue) computed from the average firing rate matrix shown in the upper panel. (C) Example neurons for the pseudo-classes with significant linear and positive quadratic components (onset and offset positive, top), significant linear and negative quadratic components (onset and offset negative, middle) and significant quadratic and non-significant linear components (on/off and duration, bottom). Color filled areas and gray lines show the TTC pattern and the corresponding fit, respectively. (D) Scatter plot showing, for each neuron, the values of the linear and quadratic coefficients of the TTC fit (same color code than C, non classified neurons in gray).

**Figure 1-figure supplement 1:**
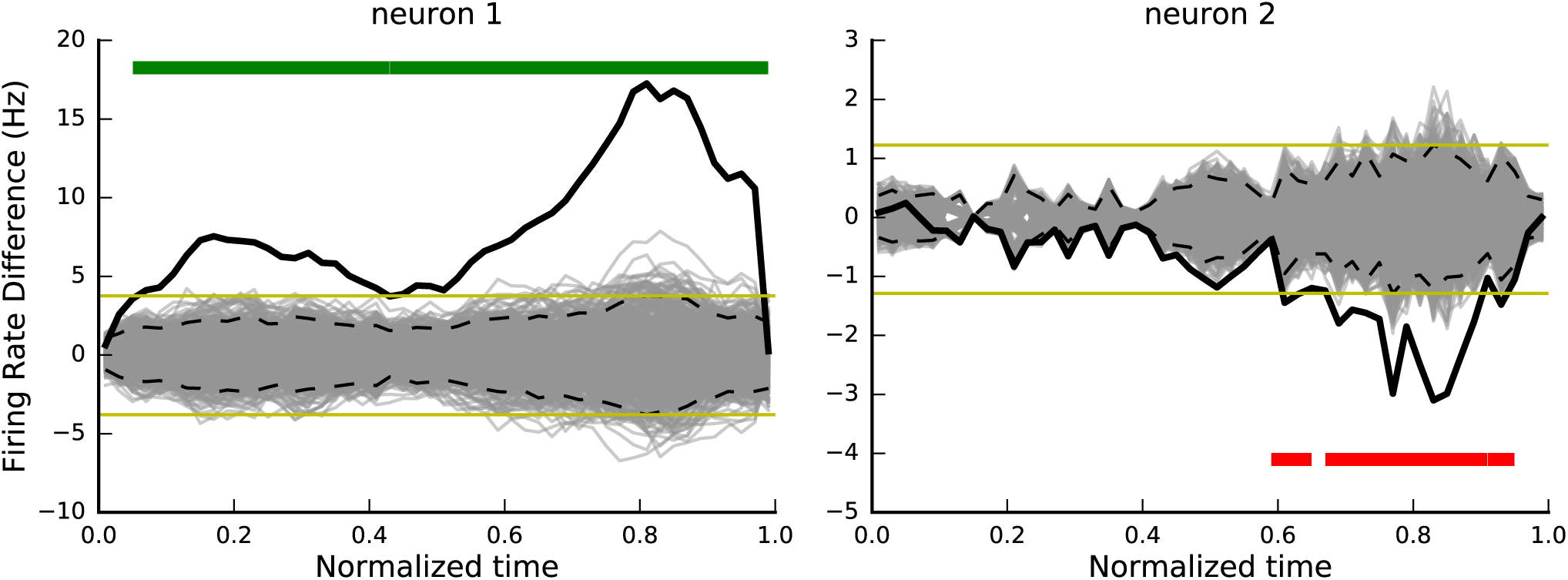
Significance of firing rate difference between trial and intertrial periods. (Left) A positively modulated neuron same than Figure 1A middle. (Right) A negatively modulated neuron same than Figure 1A bottom. Solid black and gray lines correspond, respectively, to real and surrogate (resulting from permutation procedure, 500 repetitions) differences between trial and intertrial normalized mean firing rates. Dashed black and solid yellow lines correspond to pointwise and global confidence intervals (Methods). Green and red horizontal segments correspond to proportions of significant positive and negative differences, respectively.

We first compared single neurons activities between trials and intertrials in ST animals. A majority (Methods, **Figure S1**). A majority (∼75 %) of neurons displayed a significant increase of their firing rate during trials compared to intertrials (**Figure 1A, middle panel, Figure 1-figure supplement 1**, left panel, Methods). We refer to these neurons as positively modulated. The rest of the neurons were either not task-modulated (∼3 %) or showed an increased firing rate during the intertrial (∼22 %, **Figure 1A, lower panel, Figure 1-figure supplement 1**, right panel). In the rest of the study we focused on the positively modulated neurons. After computing their mean firing rate during the motor sequence whose duration has been normalized (referred to in the rest of the manuscript as phasic tuning curve, PTC), neurons were sorted according to the phase of the sequence at which firing rate was maximal. At the population level, the modulations covered the entire motor sequence (**Figure 1B**, top). We next examined if discrete types of modulation occurring at distinct phases of the task can be extracted from the population activity. We applied a principle components analysis on the PTC of all the positively modulated neurons, which revealed that their variance can be explained by the combination of a quadratic (first principal component) and a linear (second principal component) functions (**Figure 1B**, bottom,(Panigrahi et al., 2015)). Each PTC was then fitted with a second-order polynomial function (Methods). According to the curvature and the slope of the resulting fit functions, neurons could be divided in six pseudo-classes (**Figure 1C**). Still, at the population level, the spread of the linear and quadratic coefficients revealed no clear separation between the different pseudo-classes (**Figure 1D**, mean Silhouette score of 0.17, see Methods). These results suggest that there was no discrete types of modulation occurring at specific phases of the task but rather a population response covering the entire sequence and composed of a zoo of tuning curves.

Next, we examined if the population activity is similarly modulated in HG rats, which did not learn the motor sequence. The six classes of PTC were present in both ST and HG rats (**Figure 2A-B**). However, there were more positive onset neurons (transiently active at the starting phase of the motor sequence) in HG animals than in ST animals (**Figure 2C,** proportions z-test = 2.63, P = 0.008). Conversely there was a larger proportion of negative onset neurons (active for more than half of the trial duration) in ST compared to HG animals (**Figure 2C**, proportions z-test = 4.91, P= 8.89 10^-7^). Overall, these changes were consistent with a broadening of the tuning curves after learning the motor sequence.

**Figure 2:**
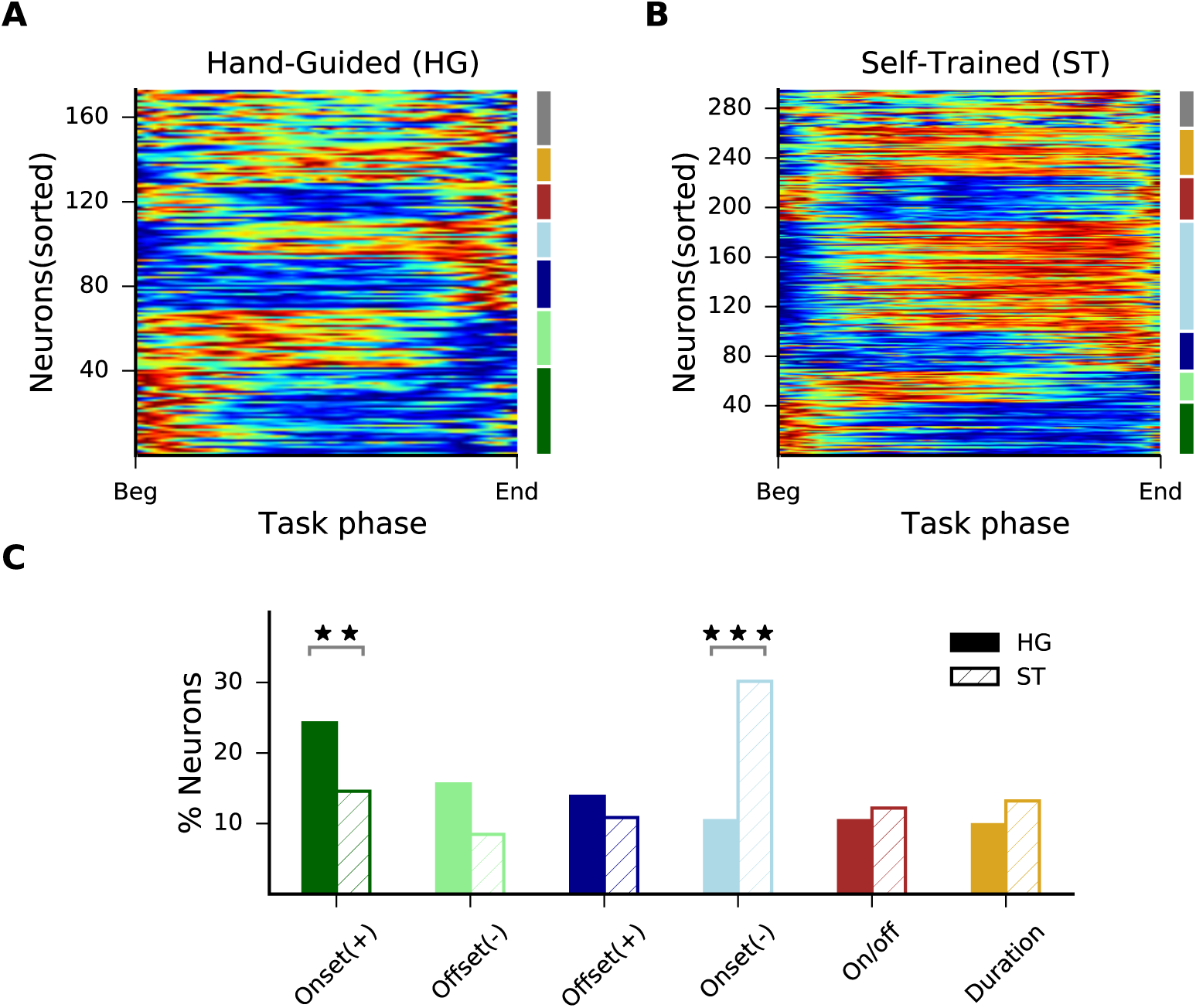
Increased proportion of broadly tuned neurons after learning. (A) Normalized average firing rates (sorted by pseudo-class, same color code as in Figure 1C) for positively modulated neurons in three HG animals. (B) Similar plot for neurons recorded in three ST animals. (C) Percentage of neurons in each class for HG (full bars) and ST (dashed bars) rats. Asterisks represent statistical significance of proportions z-test (** and *** for P *<* 0.01 and 0.001, respectively).

What could account for the broadening of the PTC observed after learning? One possibility is that striatal neurons become more sensitive to the kinematic parameters defining sequence execution, such as the position of the animal on the treadmill, running speed and acceleration. To test this possibility, we adapted a non-parametric method (Kraus et al., 2013; Saleem et al., 2013) and quantify to what extent the modulation of the average firing rate in absolute time (referred to as temporal tuning curve, TTC) can be explained by any type of relationship (i.e., not only linear or monotonic) between firing rate and the main task-relevant kinematic parameters. We generated for each neuron three kinematic tuning curves (KTC) corresponding to the average firing rate in function of the position, speed and acceleration of the animal. Then, we computed for each neuron, three model TTC computed from the KTC and the values of the rat’s kinematic parameters (position, speed and acceleration) at different times during sequence execution. We compared the real empirical TTC (**Figure 3A**) with the three model TTC (**Figure 3B**). If a given kinematic parameter was strongly influencing the firing pattern of a given neuron, then both empirical and model TTC should match. Alternatively, if the firing of that neuron was completely uncorrelated with the kinematic parameter, the model TTC should be perfectly flat. We found that the firing rate of a majority of neurons showed some degree of dependence on position, speed or acceleration (presence of non-flat model TTC) and for some neurons, the empirical and the model TTC matched perfectly (**Figure 3B**). Using a bootstrap method, we defined for each neuron and each predictive kinematic variable, a prediction quality measure (QP, QS, QA for position, speed and acceleration, respectively) as the fraction of the firing rate variance explained by each kinematic variable (i.e. a coefficient of determination). Then, we compared the influence of the different predicting kinematic variables before and after training. Noticeably, the quality of TTC prediction by running speed (QS) was better in ST compared to HG animals (**Figure 3C**) while no changes were observed for the quality of prediction by position. The influence of the acceleration on the temporal modulation was weak (QA < 0.1) in both ST and HG animals (**Figure 3B-C**). Therefore, for the rest of the study, we focused on the encoding of position and speed by striatal neurons. And for clarity purpose, neurons with high QS or QP will be referred to as speed- or position-sensitive, respectively.

**Figure 3:**
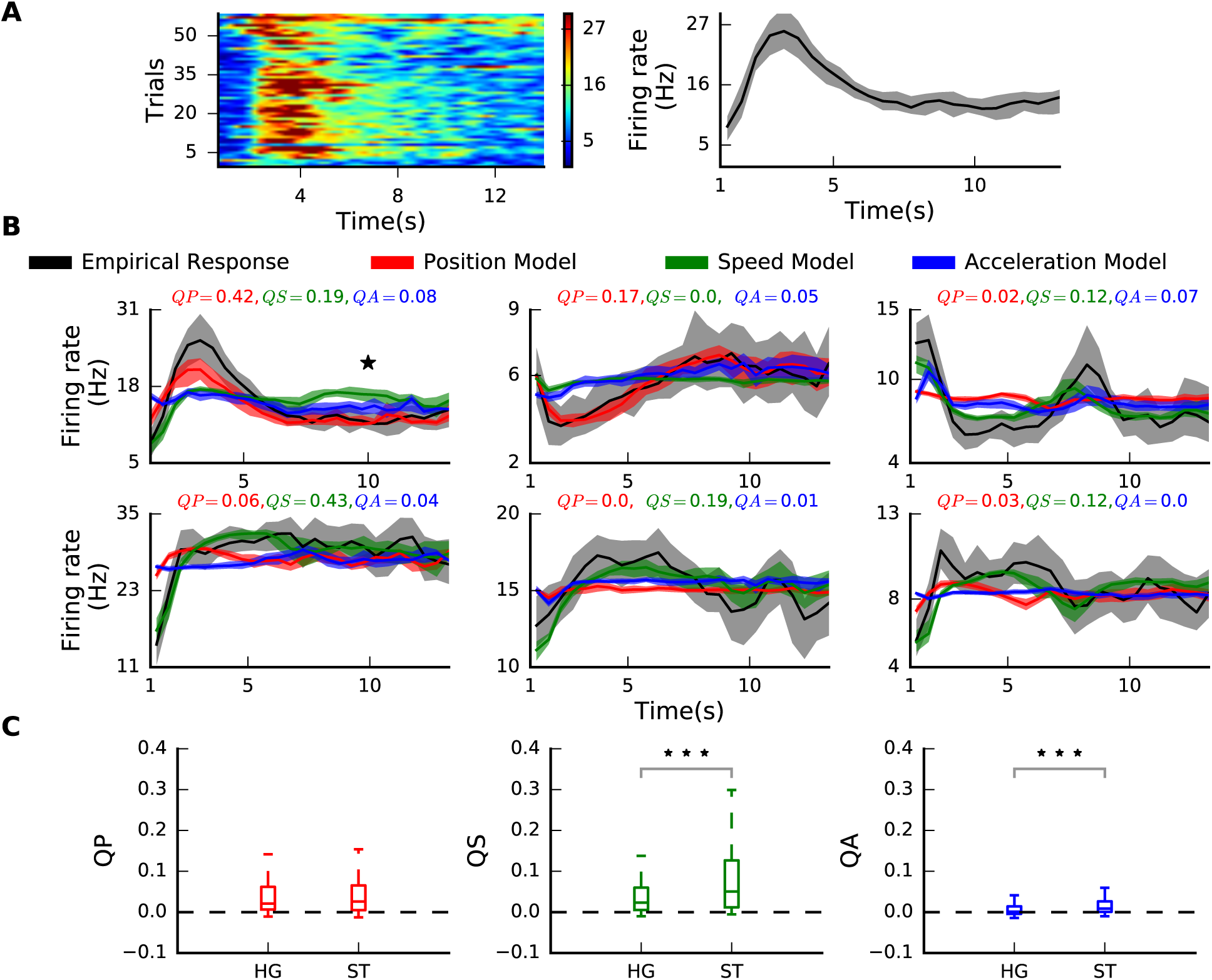
Relationship between temporal modulation of striatal neurons and task-relevant kinematic features. (A) Trial by trial firing rate (left) and TTC (right, shaded area shows +/- standard deviation) aligned to treadmill start time, for an example neuron. (B) Empirical (black lines) and predicted (colored lines) TTC for six example neurons (shaded areas shows +/- standard deviation). The values of prediction qualities for position (QP), speed (QS) and acceleration (QA) are given for each neuron. The subplot with a star corresponds to the example neuron shown in (A). (C) Box plot of QP, QS and QA (from left to right) for all neurons recorded in HG and ST animals. Asterisks represent statistical significance of Kolmogorov-Smirnov test (*** for P *<* 0.001)

Did position and speed influence independently or jointly the activity of striatal neurons? We compared the quality of TTC prediction based on each variable alone (QP or QS) versus both variables combined (QPS). A case of joint influence is shown in **Figure 4A**. For this neuron, joint influence co-occurred with separate influences of speed and position on the TTC (QPS, QP and QS are all > 0.1). In theory, joint influence (QPS > 0.1) could also coexist with weak influences of each kinematic parameter alone (QP < 0.1 and/or QS < 0.1). We found that most of the speed-sensitive neurons (QS > 0.1) did not show any sensitivity to position (QP < 0.1, **Figure 4B**). Moreover, their TTC were as well predicted by speed alone than by speed and position together (average QP/QPS > 0.7, **Figure 4C**). A similar result was obtained when considering position-sensitive neurons. Moreover, only a small proportion of neurons were jointly influenced by speed and position (**Figure 4B**, see white region of the scatter plot). Interestingly, when comparing ST and HG rats, we found strong increase in the proportion of neurons influenced by speed (QS > 0.1, **Figure 4D**, proportions z-test = 5.54, P = 2.95 10^-8^). These results indicate that position and speed were not influencing the same populations of neurons and that, after learning, the activity of striatal populations became, in average, more sensitive to the speed of the animal.

**Figure 4:**
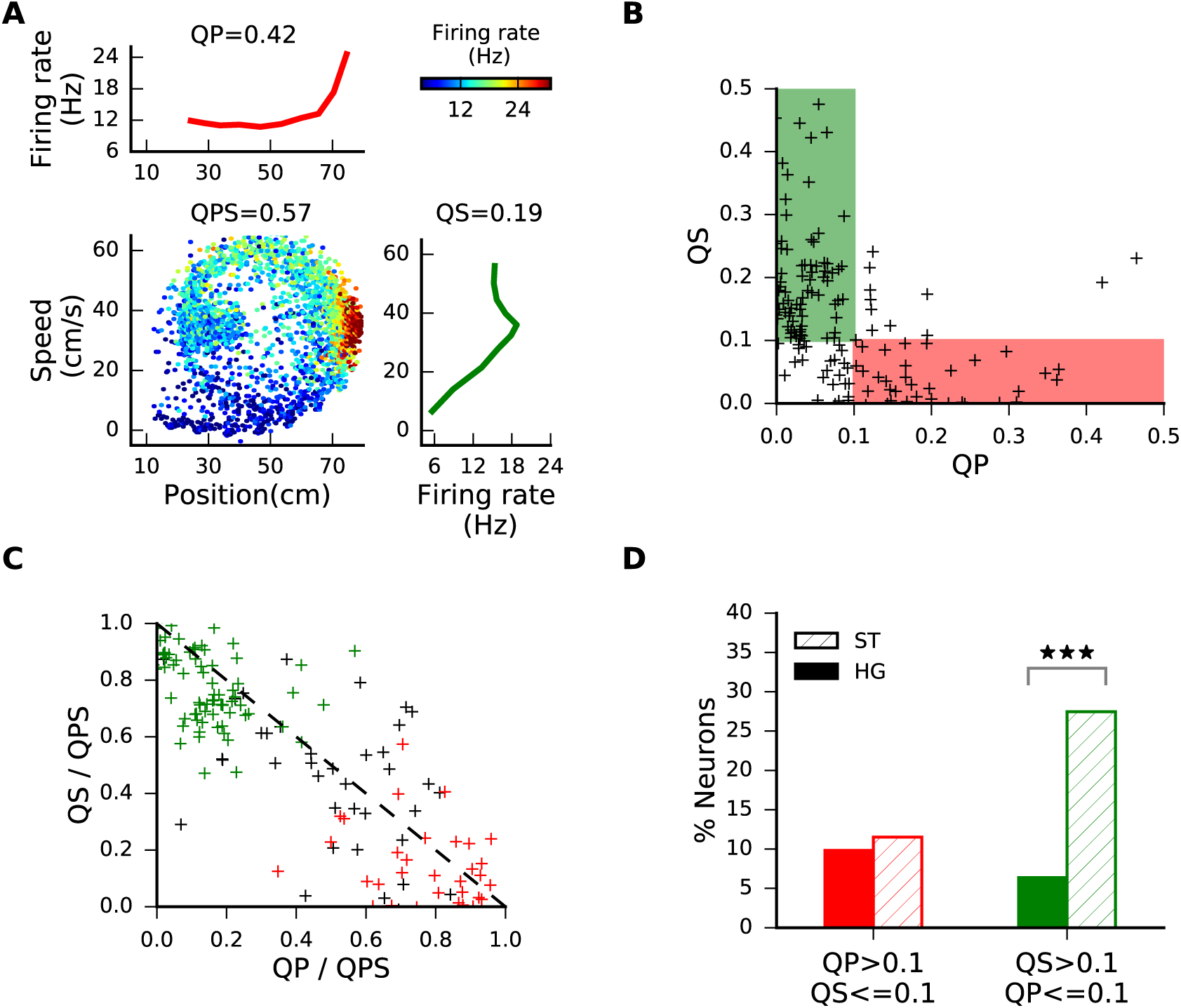
Position and running speed influence distinct neurons. (A) Mean firing rate of an example neuron versus running speed and position (bottom left), position alone (top left) and speed alone (bottom right). (B) Relative influence of speed and position on the activity of individual neurons. Only neurons from ST animals sensitive to speed and/or position are shown (QPS *>* 0.1). Green and red shaded areas correspond to neurons specifically influenced by speed and position, respectively. (C) Separate versus joint influence of speed and position on the activity of single neurons. Same color code than in B. (D) Percentage of neurons sensitive to position (QP *>* 0.1, red) and speed (QS *>* 0.1, green) in HG (full bars) and ST (dashed bars) rats. Asterisks represent statistical significance of proportions z-test (*** for P *<* 0.001)

The above analyses highlighted, on the one hand, the dependency between kinematic parameters relevant for task completion (the animals running speed and their position on the treadmill) and the spiking activity of striatal neurons and, on the other hand, the plasticity of this dependence during learning. Still, it remained to be tested if the spiking activity of striatal neurons at a given time is predictive of these kinematic parameters (where is the animal on the treadmill and at what speed it is running at that given time). To address this question we took advantage of the Bayesian inference framework (Doya et al., 2007) and developed a decoding algorithm that computed the probability of the animal being at a certain position or running at a certain speed (binned decoded features) given the firing rate of neurons at specific times (occurrences). The average probability (over several occurrences) at the bin corresponding to the rat’s real position (or speed) was used as a measure of decoding accuracy: it indicates how good the neuronal ensemble predicted the rat’s position (or speed, **Figure 5A**, Methods, (van der Meer et al., 2010)). We first showed that the decoding accuracy of single neurons was very low for position and speed (position and speed data were divided in 10 bins, i.e. chance level was 1/10, **Figure 5B**, Methods). Then, we examined the decoding performance using different sizes of neuronal ensembles (see Methods). We found that the mean decoding accuracy increased with the ensemble size (**Figure 5C**). We noted that for both, position and speed, the decoding accuracy did not seem to reach a plateau even for ensembles composed of more than 100 neurons.

**Figure 5:**
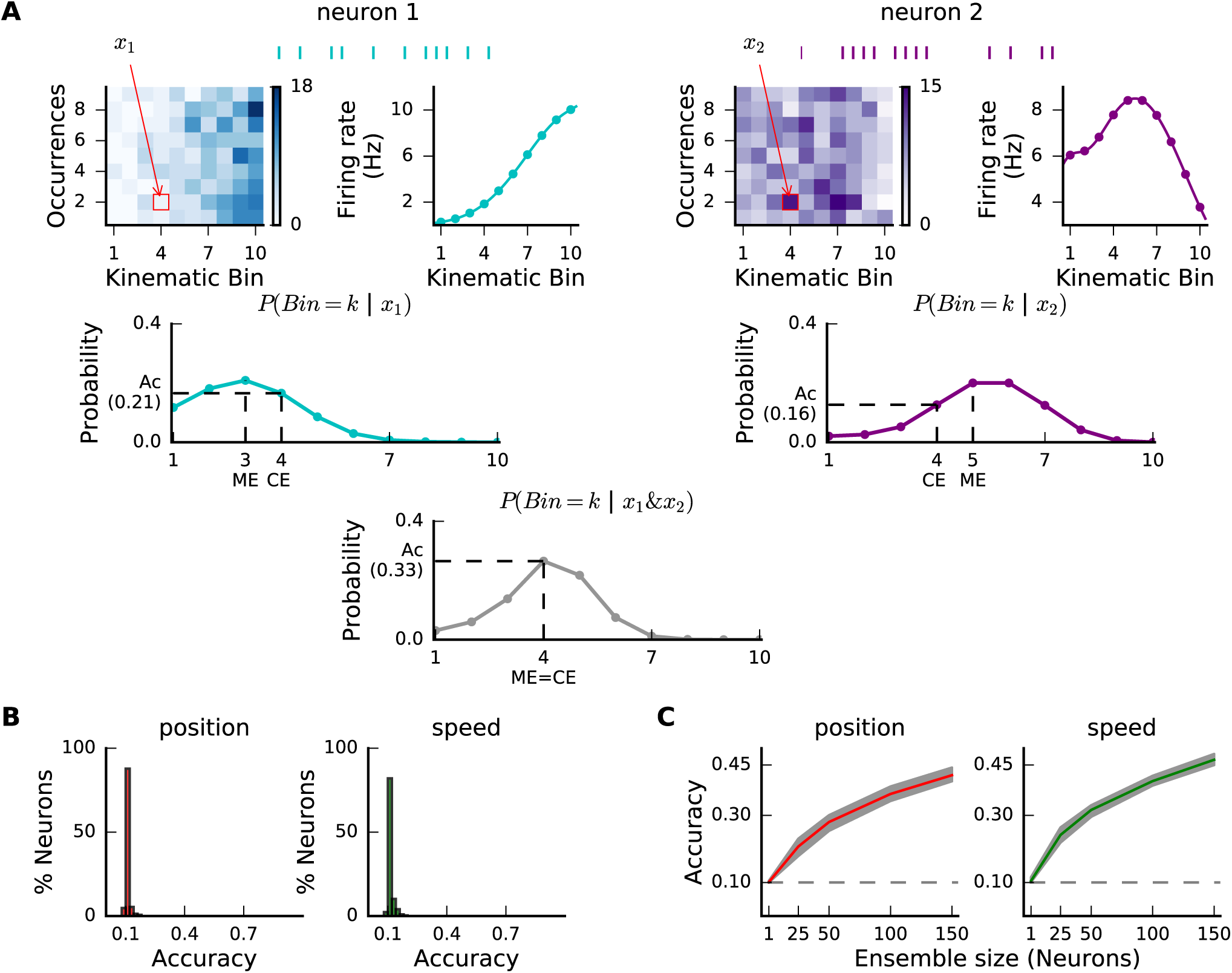
Population Bayesian decoding of position and speed. (A) Schematic representation of the decoding procedure. For each neuron, we generated a firing map corresponding to the number of spikes fired when the animal is at a given bin of position (or speed) during a given time interval (all occurrences). A position (or speed) tuning curve is generated by averaging the firing rates across occurrences. For each neuron, the posterior probability that a given firing rate (x1 for Neuron 1 and x2 for Neuron 2) corresponds to the animal being at a given position (or speed) can be computed. By assuming that neurons are independent, the posterior probability functions are combined to generate an ensemble probability function. CE: correct estimation, ME: maximum likelihood estimation. Ac: accuracy (probability of a good estimate) (B) Distribution of single neurons accuracies for position (red) and speed (green) decoding in ST animals. (C) Position and speed decoding accuracies increase with the number of neurons included in the decoding algorithm. Red and green lines correspond to the average accuracy over 100 same size subsets randomly selected, while shaded areas correspond to 25 % and 75% percentiles. Dash lines show chance level.

The relatively constant increase in mean accuracy of decoding with increasing size of neuronal ensembles (**Figure 5C**) is likely due to the fact that, to generate subsets of different sizes, neurons were randomly selected, a procedure known to level down the contribution of singles neurons (Lebedev, 2014). We hypothesized that kinematic-sensitive neurons (neurons with high QP or QS) contribute more strongly to population decoding. Indeed, we found that, for both position and speed, the gain in decoding accuracy associated with increased ensemble size was faster when incorporating first position- or speed-sensitive neurons (**Figure 6A-B**). Importantly, the top 15th percentile position- (or speed-) sensitive neurons achieved more than 70 % percent of the decoding accuracy calculated from the entire population (**Figure 6C-D**). Such level of decoding accuracy using a minority of “expert” neurons could also be reached with low sensitive neurons (lowest values of QP and QS) if more than 85 % of the population was included in the decoder (**Figure 6C-D**).

**Figure 6:**
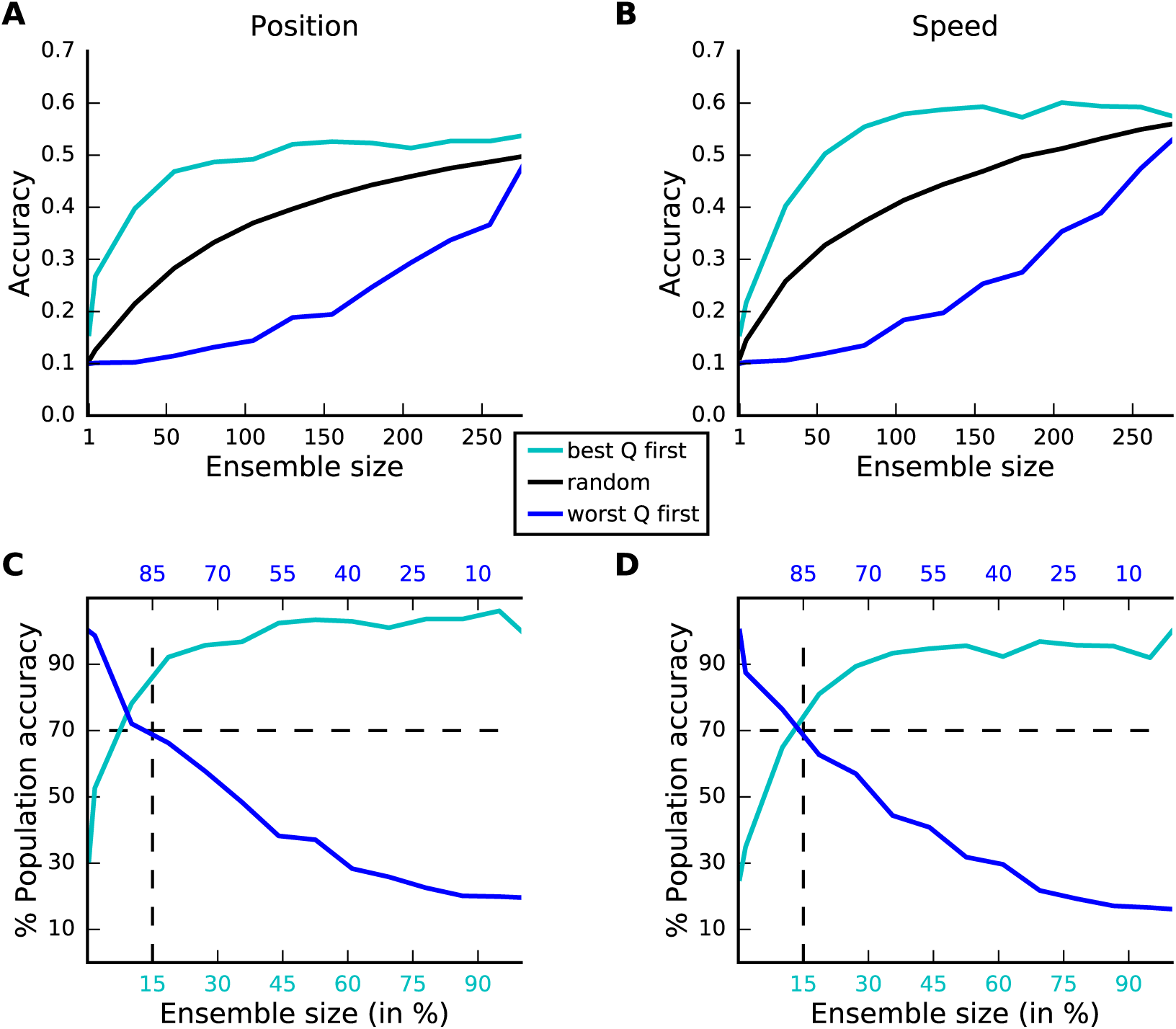
Individual neurons contribute unevenly to population decoding. (A) Accuracy of position decoding versus number of neurons used in the decoder if position-sensitive neurons (high QP) are incorporated first (light blue), if position insensitive neurons (low QP) are incorporated first (dark blue) or when neurons are incorporated randomly (black, average over 100 repetitions) (B) Similar plot as in panel (A) for speed decoding based on QS. (C) Accuracy of position decoding (in % with respect to the decoding accuracy reached using the entire population) versus ensemble size (in % with respect to the entire population size), using the neurons with best (light blue) or worst (dark blue) QP values. The position (resp., speed) decoding accuracy of the best 9 % (resp., 13%) neurons is equivalent to that of the worst 91% (resp., 87%). (D) Similar plot as in panel (C) for speed decoding based on QS values.

The above results showed that each striatal neuron serves as a specialist which encodes a specific feature of the task, as well as a generalist, which participates weakly to the population coding of other task-related features. Together with the fact that speed-sensitive and position-sensitive neurons are separate ensembles of neurons (**Figure 4**) this suggests that the striatum multiplexes kinematic information mostly at the population level. To better quantify the contribution of single neurons to population decoding, we calculated for each neuron the average difference in accuracy of position or speed decoding when this neuron was added to a group of five randomly selected neurons (Methods). We then compared, for each neurons, its contribution to position (or speed) decoding and its QP (or QS). As expected, the two measures were highly correlated (Pearson-correlation coefficients higher than 0.7) for both position and speed (**Figure 7**). Then, to optimally compare the population decoders in HG and ST animals, we used a dropping procedure that does not depend on the size of the entire population of neurons in each group and take into account the contribution of position (or speed) decoding of the individual neurons (Methods). This procedure revealed a clear gain in speed decoding accuracy in ST rats compared to HG rats (**Figure 8B**) as could be expected from the gain in the percentage of speed-sensitive neurons. However, the accuracy of position ensemble decoding was similar in ST and HG rats (**Figure 8A**). Finally, we found that the distributions of the single-neuron contributions to position and speed ensemble decoding were strongly skewed with a heavy tail (**Figure 8C-D**, empty bars), confirming the presence of a minority of position- and speed-expert neurons. Strikingly a large fraction of the neurons that contributed highly to speed encoding belonged to the group of negative onset neurons (**Figure 8D**, blue bars), whose proportion increased after learning (**Figure 2C**). This suggests that during learning a minority of neurons have become “leaders” of the striatal population and drive it to encode more accurately running speed, an essential parameter that the animals must control to perform the task properly.

**Figure 7:**
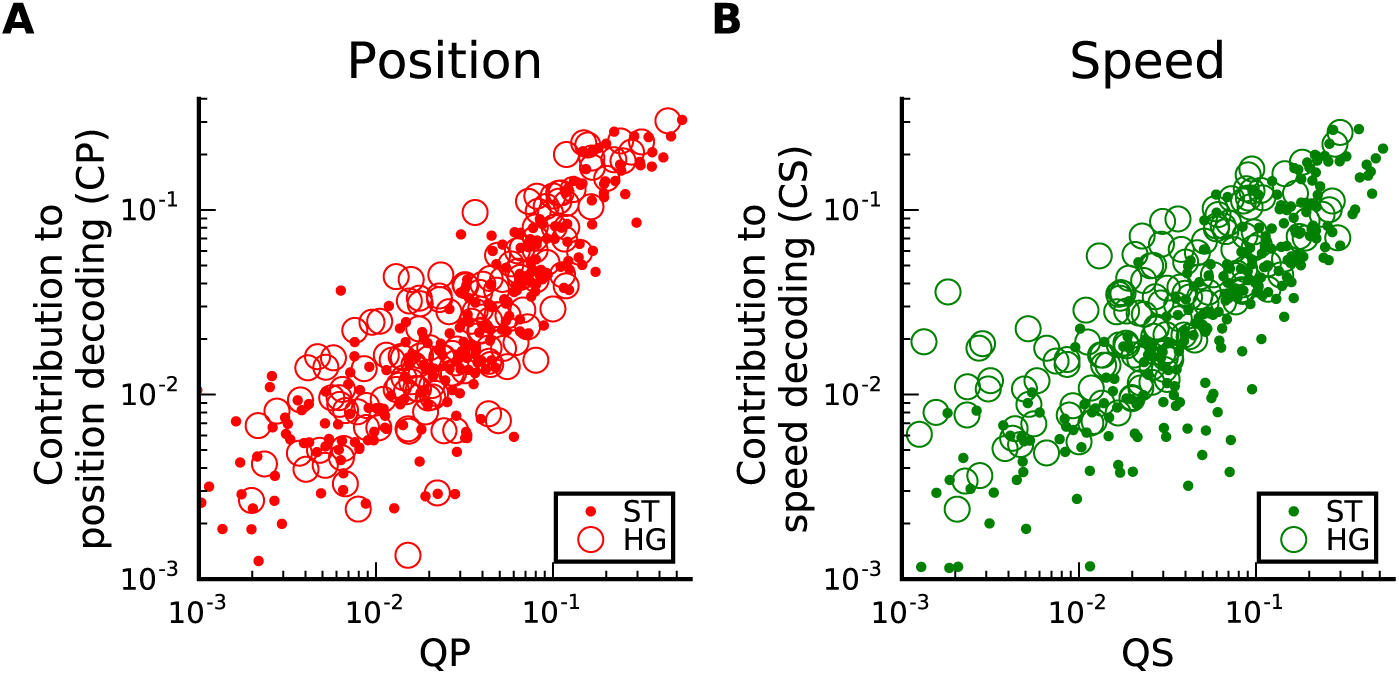
Prediction quality and decoding contribution are correlated. (A) Contribution of single neurons to position decoding (CP) versus position sensitivity of firing rate (QP) for HG (Pearson-correlation: 0.91, P *<* 0.001) and ST (Pearson-correlation: 0.90, P *<* 0.001) rats. (B) Contribution of single neurons to speed decoding (CS) versus speed sensitivity of firing rate (QS) for HG (Pearson-correlation: 0.78, P *<* 0.001) and ST (Pearson-correlation: 0.83, P *<* 0.001) rats.

**Figure 8:**
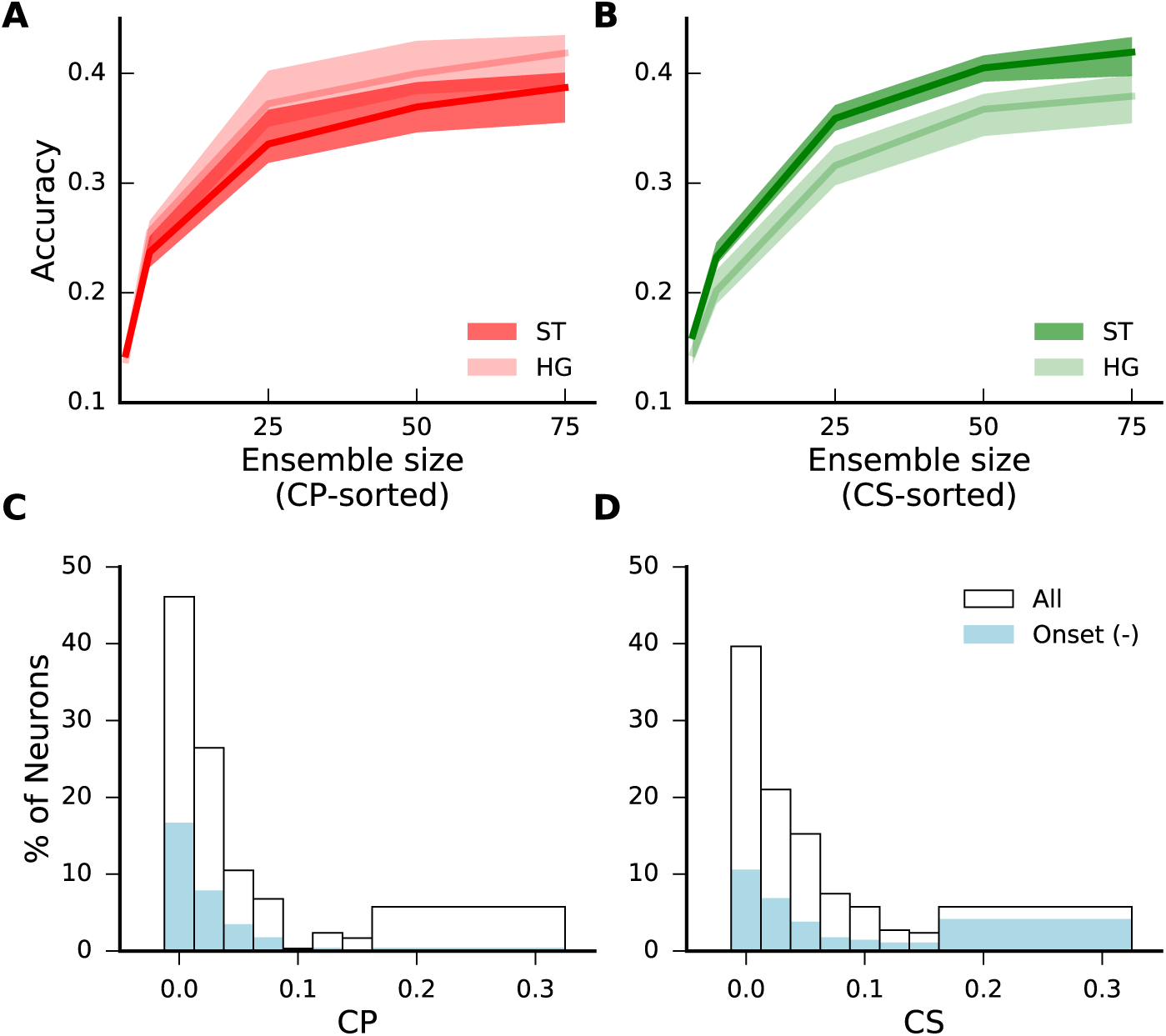
Improved running speed decoding accuracy after training. (A) Position decoding accuracy versus numbers of neurons (neurons were incorporated according to their decoding contribution). Solid lines and shaded areas correspond respectively to the average accuracy values and the 25 % to 75 % percentile intervals for neurons from ST and HG animals. (B) Same as in (A) for speed decoding accuracy with dark green for ST rats and light green for HG rats. (C) and (D) Distribution of CP and CS, respectively, for all positively modulated (empty bars) and only negative onset (light blue bars) neurons in ST animals.

## Discussion

How ensembles of neurons in the striatum encode task-relevant information has been studied in a hand-full of studies, in the context of spatial navigation (van der Meer et al., 2010) and time representation (Gouvêa et al., 2015; Mello et al., 2015; Bakhurin et al., 2016). Here, we report that ensemble of striatal neurons encoded separately two kinematic parameters (position and running speed) defining how a sustained running sequence was performed. The ensemble encoding accuracy of running speed changed after learning and was ruled by a minority of “expert” neurons specifically sensitive to this kinematic parameter. Still accurate decoding could also be obtained using the remaining “non-expert” population. Altogether our work provides, on the one hand, insight on how individual neurons contribute to striatal ensemble representation of motor sequences and, on the other hand, a flexible but robust population code to implement the recently emphasized contribution of the basal ganglia to movement vigor (Turner and Desmurget, 2010; Dudman and Krakauer, 2016).

### Heterogeneity of neuronal modulations in the striatum during motor sequence execution

Early recordings of spiking activity in the putamen of non-human primates performing well-controlled short movements have revealed neuronal responses that 1) were movement-related and covered a wide range of timing, mostly after but also before movements initiations, 2) showed context-dependent modulations to movement and 3) were sensitive to task-relevant cues or sensory stimulation (Mink, 1996). In contrast to this complexity and variability of responses to brief movements, recordings in the DLS from rodents performing learned motor sequences have suggested the existence of functionally separated classes of neurons with phasic modulations compatible with a role in the chunking and/or concatenation of action (Jog et al., 1999; Barnes et al., 2005; Jin and Costa, 2010; Jin et al., 2014). The classification of neurons in distinct functional types based on the temporal profile of their response during task performance requires unbiased methods to capture the variability of the population response and to quantify the degree of separability/overlap of the putative classes. Here, using principal component analysis, we found that the variability in population responses was well explained by a combination of a linear and quadratic functions (Panigrahi et al., 2015). The most informative aspects of the temporal response of each neuron during the task can therefore be represented in a lower two-dimensional space defined by the coefficient values of the linear and quadratic functions (which approximate the response of the neurons). This did not yield to the characterization of distinct classes of the response. Rather, our analysis revealed a diversity of heterogeneous phasic modulations occurring throughout task performance (**Figure 1B**). The DLS region in which we performed our recordings receives strong cortical and thalamic sensorimotor input that process information from different body parts (Mittler et al., 1994; Cho and West, 1997; Hintiryan et al., 2016; Hunnicutt et al., 2016). It is therefore likely that the heterogeneous responses of striatal neurons reflect the heterogeneity of time-varying sensorimotor input during task performance (Ponzi and Wickens, 2012; Barbera et al., 2016). In our population responses, the most phasic (transient) modulations of neuronal activity occurred preferentially at the beginning and end of the running sequence (**Figure 2**). Brief sensorimotor stimulation of different body parts are likely to occur during these two phases of the sequence (when animals changed posture and started or stopped to run) while more sustained sensorimotor stimulation would be associated with stable running (middle phase of the task). To sum up, high-order functions such as the chunking of actions have been attributed to the DLS on the basis of the observation of functional types of neurons active at specific phases of motor sequence but in our task such functional classes could not be isolated statistically. This obviously does not exclude that some neurons are modulated at different phases of the task. But due to the heterogeneous nature of sensorimotor information reaching the DLS it is difficult to rule out that phasic modulations reflect primarily covert sensorimotor aspects of task performance occurring regularly at certain phases of a motor sequence rather than high-order functions.

### Ensemble representation of running speed in the striatum

When we compared population responses during task performance in naive and trained animals, we found a marked increase in the proportion of neurons responding with a broad modulation of their firing rate (**Figure 2C**). We have previously argued that the difference in neuronal responses between self-trained and hand-guided conditions reflect learning-related processes rather than a difference in sensorimotor stimuli or reward expectation (Rueda-Orozco and Robbe, 2015). Based on several studies reporting linear correlations between striatal firing rates and speed of movement or locomotion (West et al., 1990; Carelli and West, 1991; Pederson et al., 1997; Tang et al., 2007; Kim et al., 2014; Panigrahi et al., 2015; Rueda-Orozco and Robbe, 2015), we examined if this increased proportion of neurons with sustained activity could be related to a change in representation of the main task-relevant kinematic parameters, the running speed and position of the animal. By representation we meant the capacity to predict speed or position from spiking activity (encoding). Indeed correlations can be statistically significant but too noisy (see examples in (Panigrahi et al., 2015; Rueda-Orozco and Robbe, 2015)) to be behavioral predictive at either single or population levels. To address this issue we adapted, on the one hand, a non-parametric method to quantify beyond linear correlations the influence of speed and position on spiking activity of single neurons (Kraus et al., 2013; Saleem et al., 2013) and, on the other hand, a Bayesian inference algorithm to test the encoding capacity of single neurons and neuronal ensemble reconstituted from different sessions. We found that single neurons hardly encoded position or running speed above chance level. However, encoding accuracy was largely improved when we considered neuronal ensembles of increasing sizes. Importantly, we found that more than 70 % of the kinematic ensemble encoding was equally performed by a minority of expert neurons (less than 15 % of the population) or the remaining less “knowledgeable” population. The idea that the activity of a minority of neurons is as determinant as the remaining majority has been suggested as a key feature of cortical network function as this allows robust and adaptative representation (Buzsáki and Mizuseki, 2014). Our study provides an original experimental support of this idea in the context of motor sequence encoding and extend its validity beyond cortical networks. Moreover, we found that, contrary to position, ensemble encoding of running speed by striatal population was more accurate after learning. Interestingly, the neurons that contributed highly to speed encoding in trained animals belonged mainly to the group of negative onset neurons, whose proportion increased after learning (**Figure 2C** and **Figure 8D**). These results suggest that the dorsolateral striatum selectively adjusted the sensitivity of single neurons to better encode running speed at the population level. In analogy with the winnerless competition theoretical framework in sensory processing (Laurent et al., 2001; Rabinovich et al., 2001), we propose that during motor learning, striatal ensembles change their task representation by tuning the activity of a minority of neurons to the kinematic parameters most relevant for task performance.

## Methods

All experimental procedures were conducted in accordance with standard ethical guidelines (European Communities Directive 86/60 - EEC) and were approved by the relevant national ethics committee (Ministère de l’enseignement supérieur et de la recherche, France, Ref 00172.01).

### Experimental methods

The behavioral and electrophysiological methods to acquire the data analyzed in the present manuscript have been described in details in a previous publication (Rueda-Orozco and Robbe, 2015). They are only briefly mentioned below.

### Running sequence task

We designed a task for rats that favors the generation of a motor sequence with fine-tuned kinematic parameters that are easily quantifiable. Specifically, we customized a motorized treadmill and trained rats to obtain rewards according to a spatiotemporal rule. Once the treadmill was turned on, animals could stop it and receive a drop of sucrose solution by entering a ‘stop area’ located at the front of the treadmill. In addition to this spatial rule, a temporal constraint was added: stopping of the treadmill was only effective if animals waited at least 7 s (goal time) before entering the stop area. If animals entered the stop area before the goal time, an error sound was played and they were forced to run for 20 s. After extensive training, rats successfully performed this task by performing a stereotyped motor sequence that could be divided in three overlapping phases: passive displacement from the front to the rear portion of the treadmill, stable running, and acceleration across the treadmill to enter the stop area. Before training, animals were first handled, then habituated to run on the treadmill at increasing speeds and finally trained to perform the task at a fixed treadmill speed (between 29 and 40 cm s-1) without the physical presence of the experimenter. The learning criterion was defined as: ≥ 72.5 % of correct trials over the last 40 trials, for ≥ 3 consecutive sessions. After habituation, some animals were first hand-guided by the experimenter (using a rectangular plexiglass plate) to perform a running trajectory highly similar to that performed by trained animals.

### Animals

Long-Evans rats (n = 5, male, 250 – 400 g) were housed in pairs (individually after surgery) in stable conditions of temperature and humidity with a constant light-dark cycle and free access to food and water. Here we analyzed the electrophysiological recordings in DLS from 5 animals: rats 1 and 4 completed at least 127 sessions before the start of recordings, rats 5, 15 and 17 were used in the hand-guided control task and were implanted immediately after the habituation period. After eight hand-guided sessions, rat 17 was also trained and recorded in the regular version of the task and reached the learning criterion after 41 sessions.

### Data acquisition and processing

Under deep isoflurane anesthesia, a 2 × 4 tetrode array (nichrome wires, 12.5-μm diameter, California Fine Wire) loaded on a NLX-9 micro-drive (Neuralynx) was implanted above the dorsal striatum through a craniotomy centered at ML = ±3.6 mm and AP = +0.6 mm from bregma. Tetrode positions in the DLS were verified using cresyl violet staining on paraformaldehyde fixed coronal brain sections (Rueda-Orozco and Robbe, 2015). Wide-band neurophysiological signals from the tetrode arrays were amplified and continuously acquired at 20 kHz. Spike sorting was performed semi-automatically using KlustaKwik (http://klustakwik.sourceforge.net) clustering software and the graphical spike sorting application Klusters (http://neurosuite.sourceforge.net). The animal position was tracked using a CCD camera (sampling rate = 60 frames s-1) positioned laterally to the treadmill and a marker attached to the left forelimb.

### Data analysis

All the analyses were performed using the Python language taking advantage of the Jupyter Notebook web interface. The behavioral and neuronal data and the notebooks used to generate the figures are available upon request to the corresponding author.

Sessions with less than 30 trials were excluded from the analyses. We considered in total 55 sessions for ST animals (23, 19 and 13 sessions in each animal) and 22 sessions for HG animals (7, 8 and 7 sessions in each animal) leading respectively to 395 and 256 isolated neurons with an average number of neurons per session of 7 (std = 3, min = 2 and max=17 neurons) for ST and 11 (std = 5, min = 4 and max = 25 neurons) for HG animals.

### Firing rates modulation during running sequence

Trials and intertrials (periods during which the treadmill was on and off, respectively) were divided in 250 ms-long non-overlapping windows. Instantaneous firing rates were obtained by smoothing the firing rate in each window (spike count / 0.25) with a Gaussian kernel filter of standard deviation 250 ms. Neurons with average instantaneous firing rate < 0.01 spikes s-1 over the recording session were not further analyzed. To normalize their variable durations, all trials and intertrials were divided in 50 non-overlapping windows. The instantaneous firing rates during individual trials and intertrials were linearly interpolated such as each neuron’s activity was described by two arrays of shape n x 50 (with n being the number of trials/intertrials in the recording session; examples of such maps are shown in **Figure 1A**, with the intertrial map duplicated for illustrative purpose). Based on these firing maps, we computed the session average firing rates vectors f _trial_ (also referred in the manuscript as phasic tuning curve, PTC) and f _intertrial_. Then, we applied a permutation test to determine whether there were significant differences between these two vectors. First, we computed the difference vector Δf = f _trial_ -flip (f _intertrial_) (**Figure 1-figure supplement 1**, solid black line). The flipping procedure is used to account for the continuity between trial and intertrial periods (difference almost null on the transition points). Then, we shuffled the labels of trial/intertrial periods, recomputed the difference vector and repeated this operation 500 times (surrogate differences, gray curves in **Figure 1-figure supplement 1**). We first defined a pointwise confidence interval (**Figure 1-figure supplement 1**, black dashed curves) as the 5^th^ and 95^th^ percentile value of the surrogate difference at each time point. Then, we defined a global confidence band as the max and min value of the pointwise confidence interval. The portions of the real firing rate difference (black curve) that lied outside the global confidence band were considered as exhibiting significant modulation of the firing rate.

### Kinematic features

The position values of the tracked left forelimb paw were projected onto the long axis of the treadmill. Animals’ speed and acceleration were estimated by computing the approximate first and second derivatives in time of its position values. Position and speed were smoothed (Gaussian kernels, sigma = 180 ms and 500 ms respectively).

### Functional classification of neurons

The phasic tuning curves (PTC) of the positively modulated neurons displayed a wide range of shapes (**Figure 1B**). We used a principal component analysis to capture the main determinants of this variability. The majority of the variance were explained by the first two components that could be respectively approximated by a quadratic and a linear functions. Thus, for each neuron, its PTC was fitted with a second order polynomial function g: g(x) = ax^2^ + bx + c, where x is normalized trial duration (between 0 and 1), a is the quadratic coefficient, b is the linear coefficient and c is the origin value. The coefficients a and b were considered significant if their p values (pa and pb, respectively) were lower than 0.05. Then, neurons were divided into 7 functional pseudo(putative)-classes according to the significance and sign of the quadratic and linear coefficients (**Table 1**). In brief, onset positive/negative neurons were mostly active/inactive at the beginning of the trials, offset positive/negative neurons were mostly active/inactive at the end of the trials, on/off neurons were mostly active at the beginning and end of the trials, duration neurons showed sustained activity. The seventh pseudo-class contained neurons with non-significant quadratic and linear coefficients.

**Table 1.** Functional classification criteria.

| pb ≤ 0.05 |  |  |  | pb > 0.05 |  |  |
| --- | --- | --- | --- | --- | --- | --- |
| a > 0 |  | a ≤ 0 |  | pa ≤ 0.05 |  | pa > 0.05 |
| g[0] ≤ g[1] | g[0] > g[1] | g[0] < g[1] | g[0] ≥ g[1] | a ≤ 0 | a > 0 | Non classified |
| Positive offset | Positive onset | Negative onset | Negative offset | On/off | Duration |  |

To evaluate if the different pseudo-classes could be considered as segregated clusters, we considered the polynomial fit of each neuron and plotted its linear coefficient versus its quadratics coefficient (as approximation of their 1^st^ and 2^nd^ principal component value). We considered that each neuron belonged to a separate cluster (one of the 7 pseudo-classes) and computed the Silhouette Coefficient score SC of all the neurons as: SC = (d’ - d) / max (d, d’) with d being the mean intra-cluster distance and d’ the mean nearest-cluster distance to the considered neuron. For each neuron the SC ranged from 1 (perfect assignment) to -1 (perfect assignment to the nearest cluster). Values near zero indicated that the neuron was equally likely to belong to its assigned cluster or to the closest one. Negative values indicated that a neuron has been assigned to the wrong cluster.

### Temporal tuning curves and kinematic tuning curves

For a given neuron, to compute empirically its temporal tuning curves (TTC), we used a time window of 500 ms. Then, we averaged across trials the values of instantaneous firing rates occurring in each time window. To build the corresponding kinematic tuning curve (KTC), we divided the kinematic range into 10 non-uniform bins (ensuring an equitable occupancy sampling) and averaged the instantaneous firing rates values corresponding to the animal kinematic value being in each bin.

### Non-parametric reconstruction of temporal tuning curves

To quantitatively evaluate the extent to which the firing rate temporal modulation of neuronal activity during task performance could be explained by a dependence between firing rate and a kinematic feature (position, speed or acceleration), we used the following procedure. For a given neuron, we first computed a KTC based on 80 % of the data (training set, see the method section “Firing rates modulation during running sequence”). Second, for each time window in the remaining 20 % of the data (test set), we used its KTC to estimate the instantaneous firing rate r’ corresponding to the actual kinematic value at that time. This gave us a prediction of the real firing rate r in that time window. The firing vectors R and R’ (corresponding to all the values of r and r’) were used to build the empirical and model TTC, respectively. Then, we used a bootstrap method to build 95 % confidence intervals around both curves. Finally, we quantified the quality score Q of predicting the temporal firing rate based on the dependence between kinematic features and single neurons firing rate (position: QP, speed: QS or acceleration: QA) as the fraction of variance explained by the model defined as:

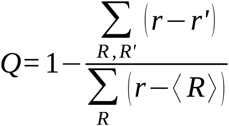

A positive quality score (max = 1 for perfect fit) meant that the model provided a better prediction of the firing rate than the constant prediction equal to the average real firing rate <R>. The same procedure was applied to compute the quality of predicting the temporal patterns based on the joint information of position and speed (QPS). The only difference was that the KTC was not anymore given by a 10 values vector but rather by a 10 x 10 array in which each value corresponded to the mean firing rate of the neuron when the animal was at a given position bin and a given speed bin at that time.

### Bayesian decoding accuracy

The decoding procedure consists in computing the probability of the animal being in a given bin of position or speed at a given time (also binned) knowing the spiking response of a single neuron (or an ensemble of single neurons) at that time (van der Meer et al., 2010). First, in a given recording session, position and speed ranges were divided into 10 non-uniform kinematic bins to ensure an equitable occupancy by the animal (all the kinematics were sampled the same amount of time (occurrence) by the animal). For each neuron, its spike train was divided in 250 ms-long non-overlapping windows and for each time window we reported its instantaneous firing rate (spike count / 0.25) in regard of its kinematic bin (e.g, the binned instantaneous position or speed of the animal at that time) to build 2D matrices as shown in **Figure 5A** (firing rate is color coded). A kinematic tuning curve (KTC) was computed by averaging the firing rate across occurrences (**Figure 5A**). For each kinematic bin k0 (k0 in [1‥10]), we applied the following procedure. For each neuron i, we took its instantaneous firing rate xi at a random occurrence of the considered kinematic bin. Second, we computed the posterior probability function p (Bin = k | xi, k from 1 to 10) of the rat being at each of the 10 kinematic bins given the KTC of the neuron, assuming a uniform prior and a Poisson distribution of the spike count variability (Hastie et al., 2009). Indeed we verified that there was no over-dispersion of the spike count (higher variability that what would be expected from a Poisson distribution) for more than 80% of the data, for both speed and position decoding in HG and ST animals (Taouali et al., 2015). The accuracy of predicting the considered bin k0 by a neuron i was given by the probability of a correct estimate CE: p (Bin = k0 | xi),(CE = 0.21 and 0.16 for illustrative neurons 1 and 2 in **Figure 5A**). To perform population decoding using an ensemble of neurons, we multiplied the individual posterior probability functions, assuming independence between the neurons included in the ensemble (which is likely due to the low level of correlation between striatal neurons (Kitano et al., 2001) and the fact that neurons were pulled from different sessions) and extracted the probability of a correct estimate from the resulting probability function (CE = 0.33 using both neuron 1 and 2 in **Figure 5A**). To evaluate the decoding performance, we applied a leave-one-out (LOO) cross validation method (Schiøtz, 2000). Then, we computed for each kinematic bin the mean probability of CE over the LOO repetitions. The final decoding accuracy of a given kinematic feature is given by averaging over the 10 considered bins the corresponding mean probabilities of correct estimation.

### Population decoding efficiency: accuracy function of sample size

We used a neuron dropping procedure (Wessberg et al., 2000) that consisted in sampling randomly 50 neuronal-ensembles of equal size from the considered neuronal population. Then, we computed the resulting average (**Figures 5C and 6C-D**) and the 50 % confidence interval (**Figure 5C**) decoding accuracy over the subsets of same size.

### Population decoding performance versus single neuron sensitivity

To investigate whether single neurons contributed unevenly to position (or speed) population decoder, we ranked all the neurons based on their QP (or QS). Then, for each considered sample size N, we selected the N best and the N worst neurons in the ranking over the entire population and computed the decoding accuracy for these two subsets (**Figure 6A-B**). We computed also for each sample size N (in percent) the decoding accuracy using the remaining (100 – N) % neurons in the population.

### Contribution of single neurons to the population decoder

The contribution of single neurons to the population decoding, was calculated for each neuron as the average difference in the accuracy of decoding resulting from including this neuron in a group of five randomly selected neurons from the considered population. Note that, apart from the consideration that single neurons may contribute reasonably to coding by such a relatively small group, i.e., leading to an accuracy above the chance level (0.1) and below the saturation level, this size was chosen arbitrarily.

### Adjusting random dropping procedure to account for uneven contribution of single neurons

To evaluate the accuracy of decoding using N neurons highly informative and at the same time randomly chosen from the population (not only the N best contributing ones over the recorded neurons), 100 neurons were selected randomly from the entire population and ranked with respect to their contribution to the decoder. Then, only the N best neurons in the ranking were used in the decoder. This procedure was repeated 50 times for each sample size to build the mean decoding accuracy curve (with 50 % confidence interval to visualize if there was a significant difference in decoding performance between HG and ST data sets, **Figure 8A-B**).

## Acknowledgments

We thank Drs Laurent Perrinet and Arnaud Malvache for critical reading of this manuscript; Members of the Robbe lab for stimulant discussions during the development of the project and Typhaine Moreau for helping to restructure the pre-processing routines in python. This work was supported by the, European Research Council (ERC-2013-CoG – 615699_NeuroKinematics, D.R.) and the Mexican Consejo Nacional de Ciencia y Tecnología (P.R.O).

## Author Contributions

WT, DR, Conception and Design, Analysis and interpretation of data, Drafting or revising the article; PRO, Acquisition of data, Drafting or revising the article

